# Thanks to repetition, dustbathing detection can be automated combining accelerometry and wavelet analysis

**DOI:** 10.1101/2023.06.24.546399

**Authors:** R.G. Fonseca, A.G. Flesia, F.C. Spanevello, M.V. de la Fuente, M.C. Bosch, R.H. Marin, L. Barberis, J.M. Kembro

## Abstract

Dustbathing is performed by many groups of birds, including Galliformes. It consists of a well-defined orderly sequence of movements. Repetitive changes in body position during dustbathing can be automatically detected through data processing of body mounted accelerometer recordings, specifically the complex Morlet continuous wavelet transform. The approach was tested in 13 adult male Japanese quail *(Coturnix japonica*) fitted with a backpack containing a triaxial accelerometer and video-recorded during at least 6h. Rhythmicity (period 25-60s) in the y-axis acceleration vector is reflected as large power values, and is associated almost exclusively to dustbathing events. Thus, by implementing a threshold value we detected events automatically with an accuracy of 80% (range 66-100%). We show potential uses for characterizing temporal dynamics (e.g. daily rhythms) of dustbathing and for the assessment of intra- and inter-individual variability over long-term studies, even within large complex environments (e.g. natural environments or breeding facilities).

**Summary statement:** We propose a method for automatically detecting dustbathing (i.e a behavior performed by many groups of birds, including Galliformes) from triaxial accerometer recoding using a wavelet technique.

## Introduction

Dustbathing is a behavior that can be found in numerous species of birds (Borchelt et al., 1973; Hogan and Boxel, 1993; Simmons, 1964; Vestergaard, 1982) and can be characterized as a precise and orderly sequence of movements, that are repeated over time, with the most characteristic movement being tossing the dust with the wings and undulating the body underneath the dust shower (Statkiewicz and Schein, 1980). Galliformes can be considered as specialists in the use of dust to bathe, given that contrary to other species they do not depend on water for purposes of integumental maintenance (Van Liere, 1991). The importance of this behavior has been documented in hens, *Gallus gallus domesticus*, that will work for an adequate substrate and in its absence will perform (sham) dustbathing even on a wire-mesh floor (Petherick et al., 1995; van Liere and Wiepkema, 1992). Moreover, deprivation of an adequate substrate also results in an increased tendency to perform dustbathing after reinstating substrate (Schein and Statkiewicz, 1983; Vestergaard, 1982). Evidence has even led to recommendations for modern legislation aiming to improve welfare to include the presence of an adequate substrate for dustbathing (AHAW, 2005) (for a review of the subject see (Olsson and Keeling, 2005)).

Dustbathing is controlled by complex interactions between internal, peripheral, and external factors (Duncan et al., 1998; Vestergaard, 1982). From diurnal observation, it has been reported that dustbathing presents circadian rhythmicity. For example, a marked peak in activity can be observed around 6 h in hens and 10 h in Japanese quail (*Coturnix japonica*) after the onset of light (Statkiewicz and Schein, 1980; Vestergaard, 1982). External stimulation, including environmental temperature, radiant heat, and light, light: dark cycles and social stimulation (Duncan et al., 1998; Hogan and Boxel, 1993; Statkiewicz and Schein, 1980) as well as the sight of a dry dusty substrate (Petherick et al., 1995), can trigger dustbathing and/or modulate its circadian rhythmicity. Thus, the study of the temporal dynamics of dustbathing can provide important insight into the bird’s adaptation to its environmental context.

To our knowledge, the only methodology available to study the dynamics of dustbathing is through visual observation of birds, mostly from video recordings. This procedure is not only time consuming but also restricts the type of studies that can be performed, limiting them to relatively small and fairly non-complex environments that can be inspected/observed easily. In this context, we propose a simple method for automating the detection of dustbathing using accelerometers and Japanese quail as an animal model. Accelerometers are sensors that can measure the acceleration vector over time in three dimensions (components x, y and z) associated with both body position and movement (i.e. static and dynamic acceleration, respectively) (Shepard et al., 2008). Accelerometers including a battery and a microprocessor can be very lightweight (<5g; https://www.technosmart.eu/axy-trek-mini/) placed within a small backpack (Fig. 1A) and attached to a bird (Buijs et al., 2018; Daigle et al., 2012; Siegford et al., 2016; Stadig et al., 2017). These accelerometers record continuously up to a week long period and can be used within natural environments given that they do not interfere with normal behavior (Pellegrini et al., 2015; Rossi, 2022; Rossi et al., 2021; Simian, 2020). Afterwards, the resulting acceleration vectors (whose components we call *a*_*x*_ *a*_*y*_ and *a*_*z*_) can be processed in order to extract behavioral information. Herein, for the detection of dust-bathing events, we propose to process the accelerometry signal using a wavelet technique. It detects rhythmicity as well as sharp changes localized in a specific time period (Flesia et al., 2022), within the obtained acceleration vectors. Specifically, we found that complex continuous wavelet transforms (cwt), such as the Morlet cwt (Flesia et al., 2022), are useful in the context of detection and characterization of dustbathing events. This is due to the rhythmic nature of dustbathing; given that it consists of a sequence of coordinated movements that are repeated consecutively over time. Also, it involves abrupt movements and changes in direction since not only pecking, scratching and dust-toss are present but also body rolls (undulation) (Statkiewicz and Schein, 1980) that could be detected in the acceleration signal.

**Figure 1.**
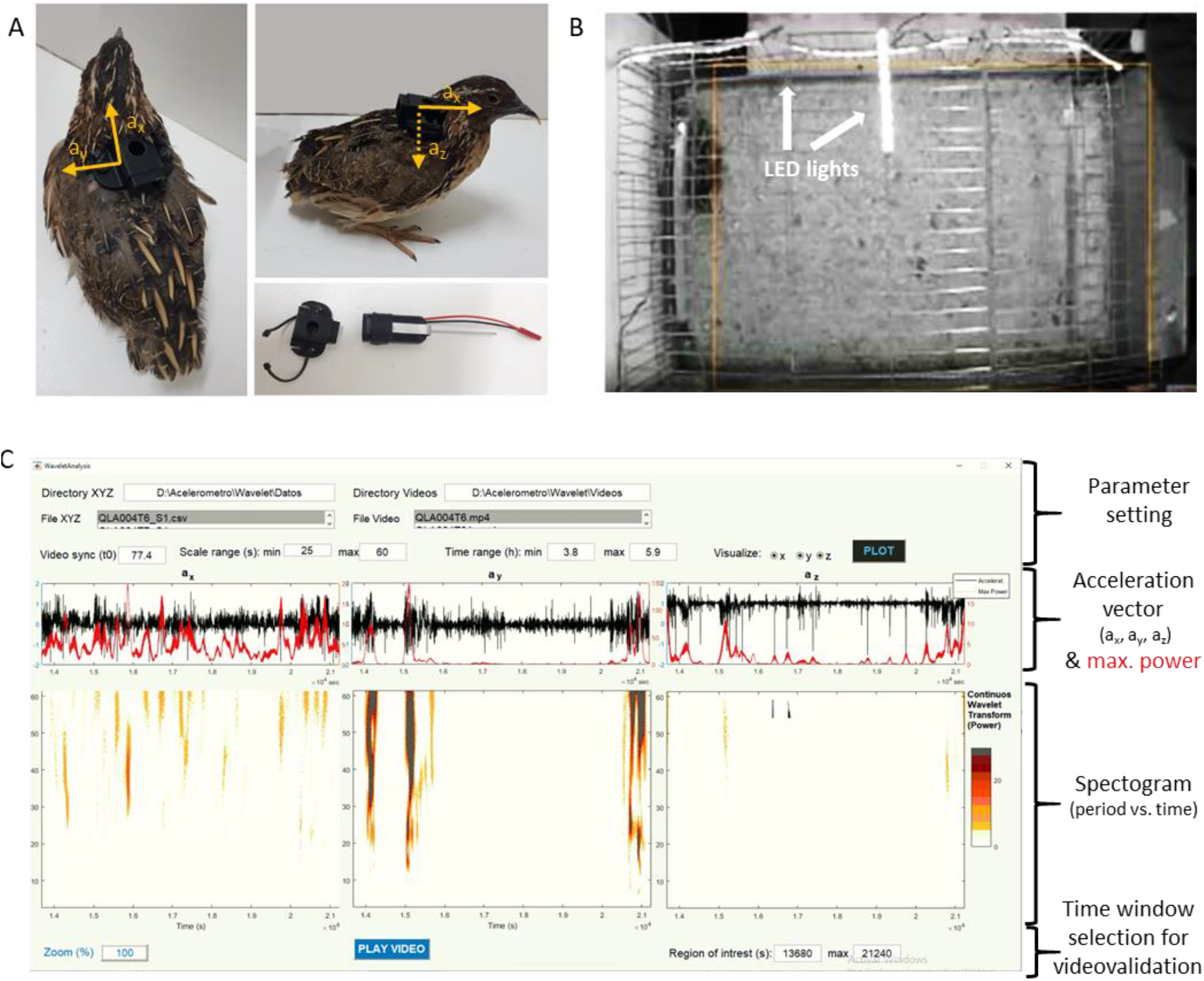
Overview of the experimental design. A) Top and side view of the male with a backpack containing an accelerometer. Arrows indicate the components of the three dimensional acceleration vector. The lower right panel shows the backpack and the accelerometer application device. B) Top view of the home box with arrows indicating the LED lights that turn on for synchronization. C) Screenshot of the MATLAB app used for validation of the wavelet method for detection of dustbathing events. The app has four sections described at the right side. The upper section/row is where users indicate the folder and file of acceleration vector data (Directory XYZ and File XYZ, respectively) and that of the corresponding video files (Directory Videos and File Video, respectively). *Parameter settings* include “Video sync” (t0)” which is the empirically established offset between video recordings and accelerometer data; “Scale range (s)” which is the range of temporal scale to be used in wavelet analysis (y-axis of the spectrogram), minimum (min) and maximum (max) values are required. In “Acceleration” (second row), the dynamics of the components of acceleration vectors are shown as black lines, from left to right, a_x_, a_y_ and a_z_ are depicted. Red lines show the maximum power (i.e squared modulus of the wavelet coefficients) values estimated for the selected range of temporal scales for each acceleration vector. The third row depicts the spectrogram showing the full range of estimated power values is observed in the section below (i.e. squared modulus of the wavelet coefficients). Yellow regions indicate large power values. The bottom section (last row) allows users to zoom into a specific time window (Region of interest (s)) and watch the corresponding video fragment by pressing the play button.

Our first aim is to validate the proposed wavelet methodology by comparing it with data obtained with the traditional visual observation of video-recordings in our animal experimental model (Japanese quail, *Coturnix japonica*) while evaluating possible technical limitations. Second, we assessed the potential of using the wavelet analysis method for the detection and characterization of dustbathing events to study the temporal dynamics of this behavior.

## Materials and Methods

Japanese quail (*Coturnix japonica*), were bred according to standard laboratory protocols described elsewhere (Caliva et al., 2019; Guzman et al., 2016). The experimental protocol was approved by the Institutional Council for the Care of Laboratory Animals (CICUAL, Comité Institucional de Cuidado de Animales de Laboratorio) of the Instituto de Investigaciones Biológicas y Tecnológicas (IIByT, UNC-CONICET). Animal care and experimental treatments also followed the Guide for the Care and Use of Laboratory Animals issued by the National Institute of Health (NIH Publications, Eighth Edition) (National Research Council, 2011). Local animal regulations were followed including the Animal Protection Law number 14346, National Administration of Drugs, Foods and Medical Devices (ANMAT) decree 6344/96, and the National Scientific and Technical Research Council (CONICET) resolution number 1047/2005.

### Animal husbandry

Eggs were collected for 10 consecutive days from adult quails from the Institute’s breeding stock. These eggs were stored in a refrigerator at 15ºC until incubation. For 17 days, the eggs were placed in an incubator/hatcher with automatic temperature and humidity controls as well as egg rotation (turning the eggs by approximately 45° every 1 h). During the first 14 days of incubation the eggs were rotated, the average temperature was maintained at 37.8°C, and the relative humidity at 65%. In the last 3 days of incubation, the eggs were transferred to trays without rotation, the temperature was reduced to 37.5°C, and the relative humidity was set at 62%. Thus, the micro-environmental conditions inside the incubator were kept optimal for embryo development.

Chicks were housed in the rearing room and distributed in white melamine rearing boxes measuring 90 × 80 × 60 cm (width, length, and height respectively) at a maximum density of 30 chicks per box. The boxes included an automatic temperature control system that was set at 37.5°C for the first week and then decreased by 3.0°C per week until room temperature (24 to 27°C) was reached in the fourth week. Quails were subjected to a daily 14L:10D h cycle (300 to 320 lx), with lights on at 6 am.

At 28 days of age, the animals were sexed according to plumage coloration. Since birds belonged to a larger experiment that evaluated aggressive and reproductive behavior, they were housed in pairs (male-female) in enriched home-cages measuring 40 × 20 × 25 cm (length × width × height). Food and water provision continued *ad libitum*.

Fourteen adult male and female pairs of at least 140 days of age were reallocated into home boxes, measuring 45 × 35 × 88 cm (length × height × width), with wire bar walls. Rice hulls covered the floor, and had an automatic waterer and a feeder.

### Experimental procedure

The experiment was designed to analyze several aspects of the quail’s behavior using accelerometry within social groups. A total of 13 groups were studied, named boxes 1-13 for future reference. However, herein we focus only on the analysis of dust bathing in males. The week prior to the experiment, a backpack was fitted onto the male, in order to promote habituation to the backpack during this period (Pellegrini et al., 2015; Rossi, 2022; Rossi et al., 2021; Simian, 2020). The backpack was 3D-printed in black plastic and had two elastic bands on their sides to be passed around the base of the animal’s wings, as shown in Figure 1A.

At least three days prior to experimentation, male-female pairs were housed in identical home boxes within an independent experimental room.

Roughly 36 h prior to the experiment, the female companion bird was placed in a separate identical home box within the experimental room, while the male remained in its home box. This physical isolation step was important since the experiment was part of a larger experiment that studied male reproductive behavior. Birds were not visually isolated from conspecifics and could visualize a pair of birds in an adjacent box.

On the morning of the experiment (between 9-11am), a TechnoSmArt@ accelerometer (9.5×15×4mm, 0.7g) was inserted into the backpack using a specially designed applicator to ensure synchronization between the acceleration time series and the two video recordings (see following section and Fig. 1A). Simultaneously, a wire bar wall partition was inserted in the center of the home box that was divided in two similar separate compartments. After placement of accelerometers, males were positioned in one of these two home box compartments. Ten minutes later, the female companion bird was placed into the adjacent compartment. After a second 10-minute period the wall partition was lifted and birds were able to interact. Video recordings began before the placement of the accelerometer and ended after a 6h period when the accelerometer was removed. As a proof of concept, the last group (box 13) was recorded during a week-long period in order to show the potential of the methodology to study dustbathing dynamics in long-term studies. Video recordings were obtained from a side camera placed 20 cm behind one of the wire bar walls, and a second camera suspended 1 m above the box. Both cameras were connected by a closed circuit to a computer.

Three time series for the components of the acceleration vector (a_x_, a_y_ and a_z_) were obtained for each male (Fig. 1A). A sampling frequency of 25 Hz (i.e. 25 data points per second) was set up based on previous studies that show the possibility of high-speed transitions between behaviors in this species (Simian, 2020). Thus, for each animal at least 540,000 time points were obtained over the 6h experimental period.

### Synchronization between the acceleration time series and the two video recordings

Insertion of the TechnoSmArt@ accelerometer into the backpack was performed with a specially designed applicator (Fig. 1A) that functioned in the following manner: the lever of the applicator is pulled and securely locked in place, allowing for the placement of the accelerometer inside the applicator and its attachment to the backpack. Once the accelerometer is secured, the lever is released, and a spring mechanism propels the accelerometer into the backpack, resulting in a peak in the acceleration record. Simultaneously, two strings of LED lights are illuminated within the home box (Fig. 1B), and that visual signal is captured by both cameras, thereby enabling the synchronization of the three recordings. When inserted into the backpack, the accelerometer remains on the dorsum above the scapular area.

### Video analysis

Video recordings were first scanned by an observer trained to detect dustbathing events. The time of beginning and end of each dustbathing event were recorded from video-recordings. By strict definition, dustbathing can be characterized as a precise and orderly sequence of movements (Statkiewicz and Schein, 1980) consisting of (a) pecking alternately from side to side with a closed beak; (b) scratching (one foot at a time) while sitting or squatting; (c) tossing the dust with the wings and undulating the body underneath the dust shower; and (d) occasionally rubbing head and/or body parts in the dust. Movements b and c, and sometimes a and d, are repeated a variable number of times. Since the initial pecking and scratching also appear in different behavioral contexts, the dust-toss and body-roll (undulation) were considered as firm indicators of dustbathing (Statkiewicz and Schein, 1980). Pauses longer than 5s, movement away from the site of dustbathing (usually preceded by standing and shaking), and/or the performance of another behavior, marked the end of each dustbathing event.

### Wavelet analysis of accelerometer recordings

Detection and characterization of dustbathing events were performed using wavelet analysis. Wavelets have the potential to describe the data without making any assumptions about trends of the component signals, and are resilient to noise (see review in (Flesia et al., 2022)). Also, information regarding changes in temporal dynamics over the length of the experiment at different time scales is quantifiable.

Wavelet analysis consists in comparing the signal (i.e. acceleration) to a function (i.e. the wavelet), at various scales and positions. Herein a continuous wavelet transform based on the complex Morlet mother wavelet was used. The selection of the wavelet was based on its capability of detecting oscillations even within noisy, non-stationary time series (Flesia et al., 2022; Guzman et al., 2017). Since a complex wavelet has both real and imaginary coefficients, the power of an oscillation can be estimated as the squared modulus of these coefficients. The power represents the strength of the oscillation at a given time scale (i.e. frequency) (Flesia et al., 2022; Guzman et al., 2017). The spectrogram represents the power as a heat map or a contour plot, setting in the vertical axis the time scale or frequency, and in the horizontal axis the time associated with the original time series. With a custom made, publicly available MATLAB application (Fonseca et al., 2023), the time series for the three acceleration components (a_x_,a_y_ and a_z_) were first studied through visual inspection of the three corresponding spectrograms. For spectrograms, power was estimated for temporal scales between 25 and 60 seconds (see below example in Fig. 1C). The maximum value of the power for each time point within the established temporal scales (i.e. between 25 and 60 seconds) was estimated.

For exploration and later corroboration of threshold values (see following section) the same customized MATLAB application (Fonseca et al., 2023) was used (Fig. 1C). This app allows users to simultaneously visualize the three acceleration components (a_x_, a_y_ and a_z,_ Fig. 1C, black lines in the panels of the second section) as function of time, their corresponding wavelet analysis shown as both, the maximum power values (Fig. 1C, red line in the panels of the second section) and the complete spectrogram (Fig. 1C, heat maps in the third section) within a specified parametric range (Fig. 1C, Scale range presented in upper section of parameter settings). Users can also zoom-in on a specific time window and then play the corresponding video fragment (Fig. 1C bottom buttons). A video showing the use of the app is presented in Video A1.

### Data analysis and classification of acceleration data

As we may observe in Figure 1C, the events marked as dustbathing consists of several concatenated sharp and rhythmic movements producing a steep increase in the size of the wavelet coefficients, depicted in the third row by color-scaled traces, in a consecutive number of scales, in a sharp period of time. This peculiar pattern is consistent with those detected automatically with thresholding techniques. In fact, this type of segmentation procedure was already suggested by Collins et al. (2015) where they propose a general method that centers on examining the shape of histograms of basic metrics derived from acceleration data to objectively determine threshold values by which to separate behaviors. Herein, we use as a metric for classification the maximum power (i.e. squared modulus of the wavelet coefficients) estimated from the acceleration vectors. Also, we compare two histograms of segmented data (i.e. with and without dustbathing, see below) to determine the appropriate threshold. This segmentation step was necessary given the disproportionately low total number of data points corresponding to dustbathing in comparison to those without dustbathing (approximately 140,000 vs 8,500,000 data points). The adapted flowchart of the process for assigning behaviors to accelerometry data (Collins et al., 2015) is shown in Figure A1. As shown in the flowchart, the process begins with a fully annotated database of synchronized video and accelerometer data (top left of Fig. A1). As stated previously, from the experiment two complementary databases were obtained for all males studied: i) a list of the time of initiation and finalization of each dustbathing event recorded from the observation of video-recording, and ii) the maximum power obtained from wavelet analysis of each of the acceleration vectors. By combining both databases, the values of the vector of the maximum power values were segmented into fragments corresponding to time periods with dustbathing events and inter-event periods, dedicated to other behaviors observable in video-recording (Fig. A2). Data from all animals during the 6h of testing was pooled together. Violin plots showing the distributions of the maximum power value for each of the pool of segments (i.e. other and dustbathing events) were performed in Python using the seaborn function, seaborn.violinplot (Waskom, 2021). Probability distributions were also performed for the maximum power value for each of the pool of segments (i.e. other and dustbathing events) in MATLAB using the histogram function.

### Accuracy and confusion matrix of classification criteria of dustbathing events

The performance of the classification criteria was assessed by estimating a confusion matrix, and associated parameters. When evaluating a standard classification model, predictions fall into four categories: true positives, false positives, true negatives, and false negatives. Dustbathing events and inter-event (i.e. time between dustbathing events where the animal is performing other behaviors) periods detected from visual observation of video-recordings were considered as ground truth, and dustbathing events and inter-event periods detected automatically from wavelet analysis of accelerometer recordings were considered as a prediction. Precision is computed with the following definitions:

- A **true positive** is observed when a prediction-target dustbathing event pair has an overlap larger than 50%.
- A **false positive** indicates a predicted dust bathing event has no associated ground truth dustbathing event or a prediction-target dustbathing event pair has an overlap smaller than 50%.
- A **false negative** indicates a ground truth dust bathing event has no associated predicted dustbathing.
- A **true negative** indicates a ground truth associated with a predicted without-dustbathing event.

Accuracy was estimated as the sum of true-positives and true-negatives divided by the total number of events, sensitivity as the number of true-positives divided by the sum of true positives (TP) and false negative (FN), TP/(TP+FN), and specificity as the number of true negatives (TN) divided by the sum of false positives (FP) and true negative, TN/(FP+TN).

## Results and Discussion

In order to be able to implement the use of wavelet analysis of acceleration vectors for the detection and characterization of dustbathing events a necessary step is to establish a standardized criterion to classify acceleration data. We first started by comparing the probability distribution of the maximum power estimated in temporal windows (i.e. segments) between “other” and “dustbathing” events as assessed from video-recordings. The violin plots shown in Figure 2A (see also histograms in Fig. A3) shows that the largest difference between distributions is observed in regard to the y-axis component of the acceleration. There are just 1.5% (119,118 out of 8,449,609) of the data points in segments dedicated to other behaviors against 77% (136,566 out of 178,350) of those with dustbathing present power values above 20 (vertical blue line, Fig. 2A). Thus, for detection of dustbathing events, focus must be placed exclusively on the maximum power estimated with wavelet analysis from the y-axis acceleration vector. In order to avoid short transitory periods of low power values, a dust bathing event was considered finalized when, for the following 5 seconds, no values above the threshold were observed (Fig. 2B).

**Figure 2.**
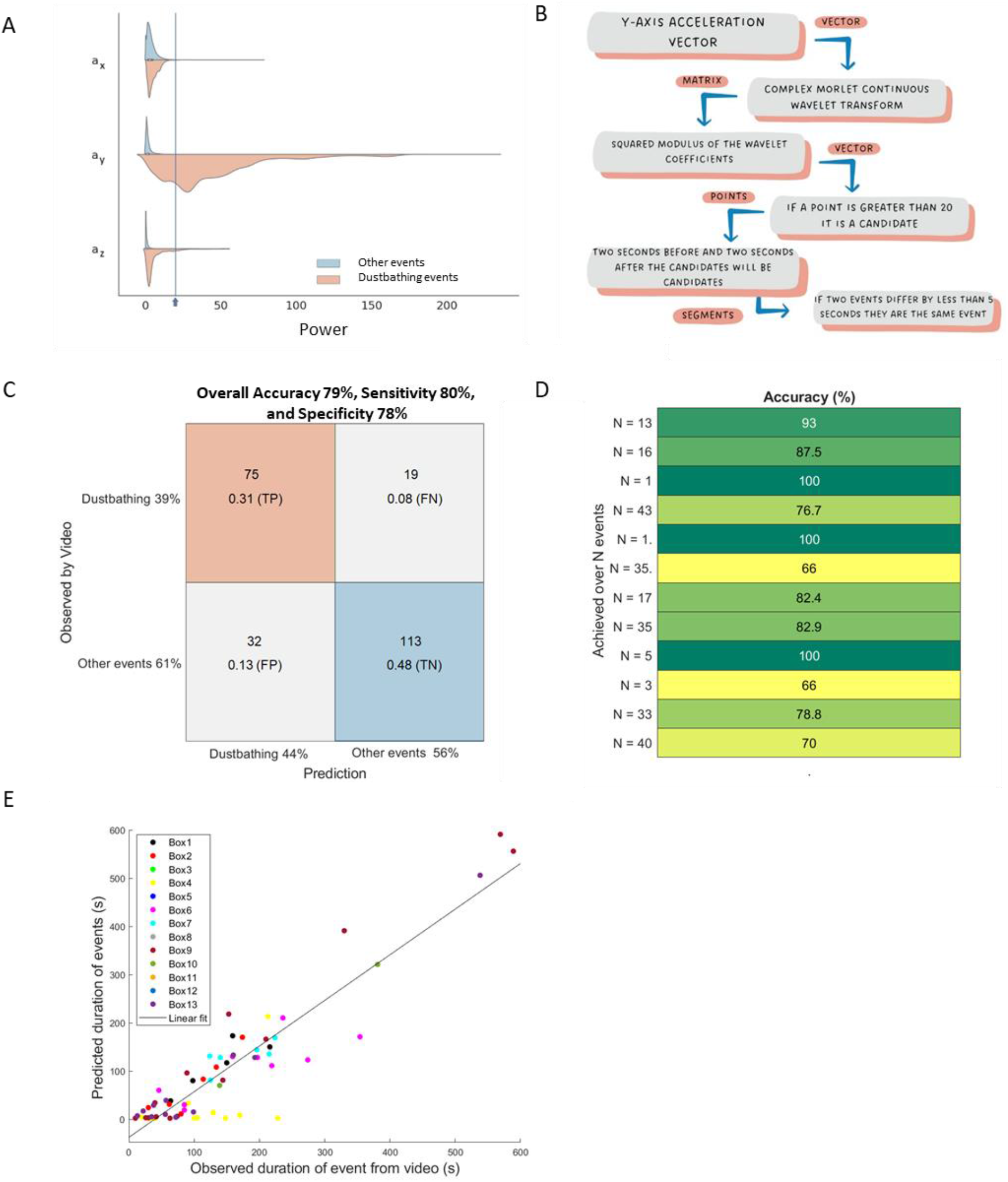
Data analysis and accuracy of dust bathing detection and characterization with wavelets. Violin plot of the distribution of values of the maximum power (i.e. squared modulus of the wavelet coefficient) estimated with wavelet analysis from x-, y- and z-axis acceleration vectors (a_x_, a_y_ and a_z_) in time windows without (light blue) and with (red) observable dustbathing events in video-recordings. Classification tree. C) Overall confusion matrix of detection of dust bathing estimating considering 12 different males (one male did not perform dust bathing during testing). The number of events in each class and the proportion of true-positives (TP), false-positive (FP), true-negative (TN) and false-negative (FN) rates are shown in squares. The percent of events from each class is shown next to the label. D) Variations in accuracy between males in the detections of dustbathing. E) Relationship between the estimation of duration of dustbathing with wavelets and the duration registered by visual observation of video recordings. The combination of symbols are colors that represent each individual.

An example of dust bathing detection based on wavelet analysis is shown in Figures A4 and A5. Zoom in on the fragment where dust bathing was both observed in video-recording (blue boxes, Fig. A5) and automatically detected using our wavelet technique (red boxes, Fig. A5). It is noticeable that the y-axis acceleration vector changes regularly between positive and negative values of acceleration. These stable values in the acceleration vector can be associated with the animal’s body position, i.e. static acceleration (white arrows in photographs Fig. A5 and Video A2). The accelerometer on the bird’s back is inclined in regard to the y-axis when the animal is laying on its right or left side during dust bathing. This inclination of the accelerometer with respect to the earth’s gravitational field is thus observable in the y-axis acceleration component and provides a measure of the body angle of the animal (see review in Shepard et al., 2008). Interestingly, our wavelet analysis shows that the bird repeats these shifts in body position periodically every 25 to 60 seconds, as denoted by the large power values for this range of temporal scales (Fig. 1C, 2A). In other words, the repetition of the sequence of movements associated with dust bathing is observable as a rhythm in the y component of the acceleration vector with a period that varies between 25-60s. Video A2 provides a visual example comparing video-recordings of behavior, acceleration vectors and the maximum power estimated with wavelet analysis of acceleration vectors.

The confusion matrix of the wavelet method for the detection of dust bathing events is presented in Figure 2C. Overall accuracy is 79%, with the probability of a dustbathing event going undetected around 8% (Fig. 2C). Sensitivity, which measures how apt the model is to detect events in the positive class (i.e. dustbathing events), is 80%, indicating the percent of actual dustbathing events observed in video-recordings that were correctly predicted as dustbathing events. Specificity, a measure of how exact is the assignment to the positive class to a segment of data is 78%. Hence, less than 22 % of all general segments are predicted incorrectly as dustbathing events.

Interestingly, accuracy varied between 66 and 100% depending on the male, independently of the number of dustbathing events performed by the male (Fig. 2C). Visual inspection of misclassifications cases (Table A1) showed that false-positives were associated with sequences where birds walked or sat, then stood up and performed grooming, while false-negatives were predominantly associated with incomplete short dustbathing events. It should be noted that accuracy was assessed based on the strict criteria of event detection in Figure 2. However, if individual data points are used instead of events, an accuracy of 97%, a sensitivity of 60%, and a specificity of 99% is observed (Fig. A6).

The relationship between the estimated duration of dust bathing events with the wavelet method and the recorded event duration from observations of video recordings is shown in Figure 2E. A linear relationship is observed with a slope of 0.95 and a determination coefficient (R^2^) of 0.85.

We end by showing examples of the potential of our methodology for studying dust bathing behavior in Figure 3. In panel A, the number of dust bathing events detected using the wavelet method was plotted as a function of time, with each color representing the dust bathing dynamics of different males within their home box. More dust bathing occurred between 8 and 10 h after lights were turned on. This is consistent with previous studies that showed daily dust bathing rhythms in Japanese quail peaked 10h after the onset of light when on a 14L:10D cycle (Statkiewicz and Schein, 1980) as the one used herein. When the number of events observed over time is divided by the total number of events performed by the male, a similar pattern is observed (Fig. 6B). However, inter-individual variability is evident in both A and B, which is consistent with the high coefficient of variation (41%) shown by Statkiewicz and Schein (1980) in regard to the time spent dust bathing at the peak time. An actogram of the dust bathing behavior of one male (box 13) over a 6-day period is shown in Figure 3C. As expected, dustbathing was detected predominantly during the daytime however, the daily pattern varies substantially. Statkiewicz and Schein (1980) also showed considerable fluctuation in the time spent dust bathing among sample days in the same individual. For example, they showed data from a male that spent 43 minutes’ dust-bathing one day and only 7 minutes two days later, representing a 6-fold difference between these days under constant environmental conditions (Statkiewicz and Schein, 1980). Moreover, it is apparent that dustbathing can occur throughout the daylight period and its temporal organization may go beyond daily rhythms (Fig. 3D) as has been observed for other quail behaviors (Guzman et al., 2017; Kembro et al., 2013; Kembro et al., 2023; Kembro et al., 2008). These results not only further show that the methodology proposed allows for automatic recording of dust bathing time series using accelerometry, but also its potential to study the temporal dynamics of dustbathing over long periods of time.

**Figure 3.**
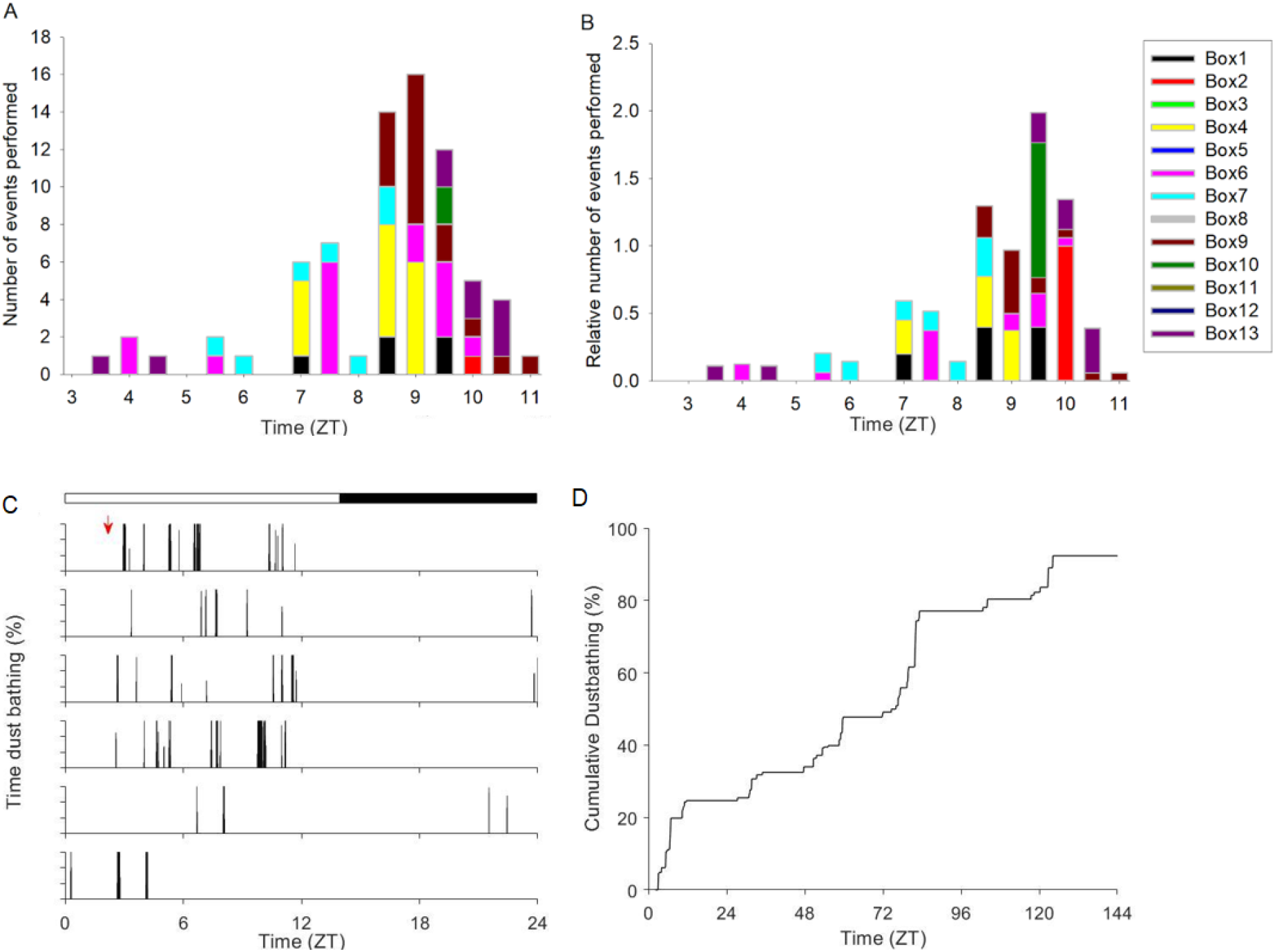
Temporal distribution of dustbathing during the light period of a 14L:10D regime. A) Number of dust bathing events as a function of time. B) The relative number of dust baths, recorded in intervals of 30 min, is estimated by dividing the number of observed events in the time interval by the total number of events performed by the male. The color code represents each male. C,D) Temporal analysis of the long-term recordings obtained from one male (box 13). In actograms (panel C) each row represents one of the 6 days recorded, and the height of the vertical bars indicate the percent of time spent dust bathing in 6 min intervals. The white/black top bar indicates the 14L:10D cycle. The red arrow indicates the initiation of testing. Cumulative dustbathing (panel D) pattern over the 6-day period. ZT: Zeitgeber time, where 0 represents the time the light is turned on.

The proposed technological approach, based on combining accelerometer recordings with wavelet analysis, will open the field to the study of the temporal dynamics of dustbathing, including daily rhythms, as well as modulatory effects within natural and complex environments. This is important since this field is currently plagued with technical limitations and time-consuming approaches of detecting dustbathing by visually observing video-recordings. Hence, to date, few studies have been designed to understand long-term patterns of dustbathing within social groups, or even daily rhythms in different experimental contexts. Also, the proposed technological approach can be used to further explore the relationship between dustbathing and poultry welfare by expanding on the premises proposed by Olson and Keeling (2005). Due to the ease of obtaining dustbathing from accelerometer recording, future research can focus on the two sides of the welfare issue. On one side, by studying both the importance of performing the behavior *per se* and thus the direct effect on welfare if the behavior is frustrated. On the other side, the possible secondary consequences of not being able to dust bath, such as deterioration of the plumage, and the effects that these may have on welfare (Olsson and Keeling, 2005). Moreover, by using accelerometer recordings linked with wavelet analysis we will be able to assess how different substrates, housing conditions, and social relations not only modulate the amount of time performing dust baths but also their temporal distribution over time and potential changes in daily patterns.

## Data availability

Dust bathing events obtained from video-recordings as well as acceleration vectors recorded with accelerometer will be available upon publication on figshare (de la Fuente et al., 2023).

## Appendices

**Figure A1.**
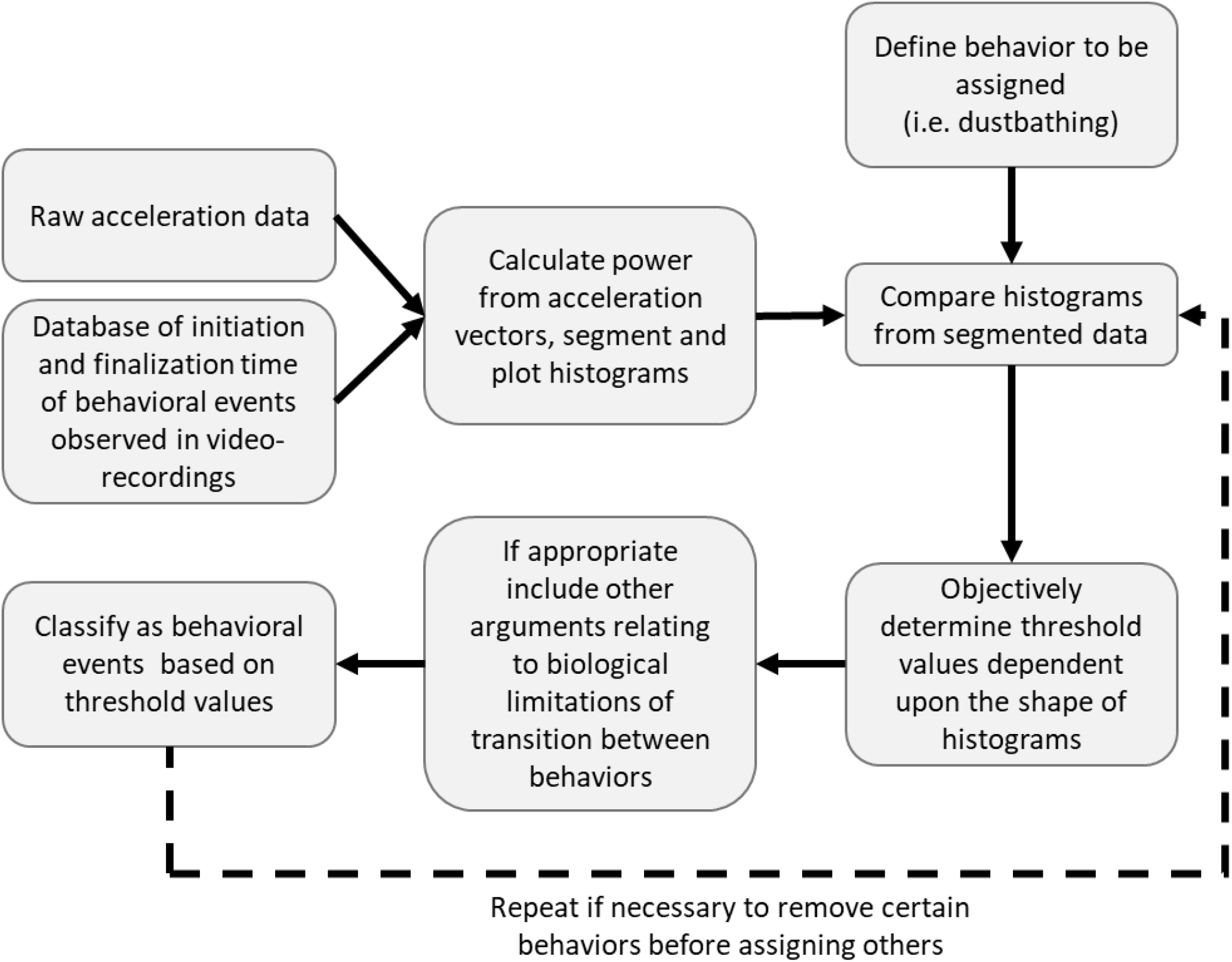
Flowchart of the process for assigning behaviors to accelerometry data (adapted from Collins et al (2015), doi: 10.1002/ece3.1660). The process begins with databases of raw acceleration data and recorded behavioral events obtained from the corresponding video-recordings (upper left corner), and ideally ends when the power values estimated from acceleration data is classified as behavioral events (i.e. dustbathing events) based on threshold values.

**Figure A2.**
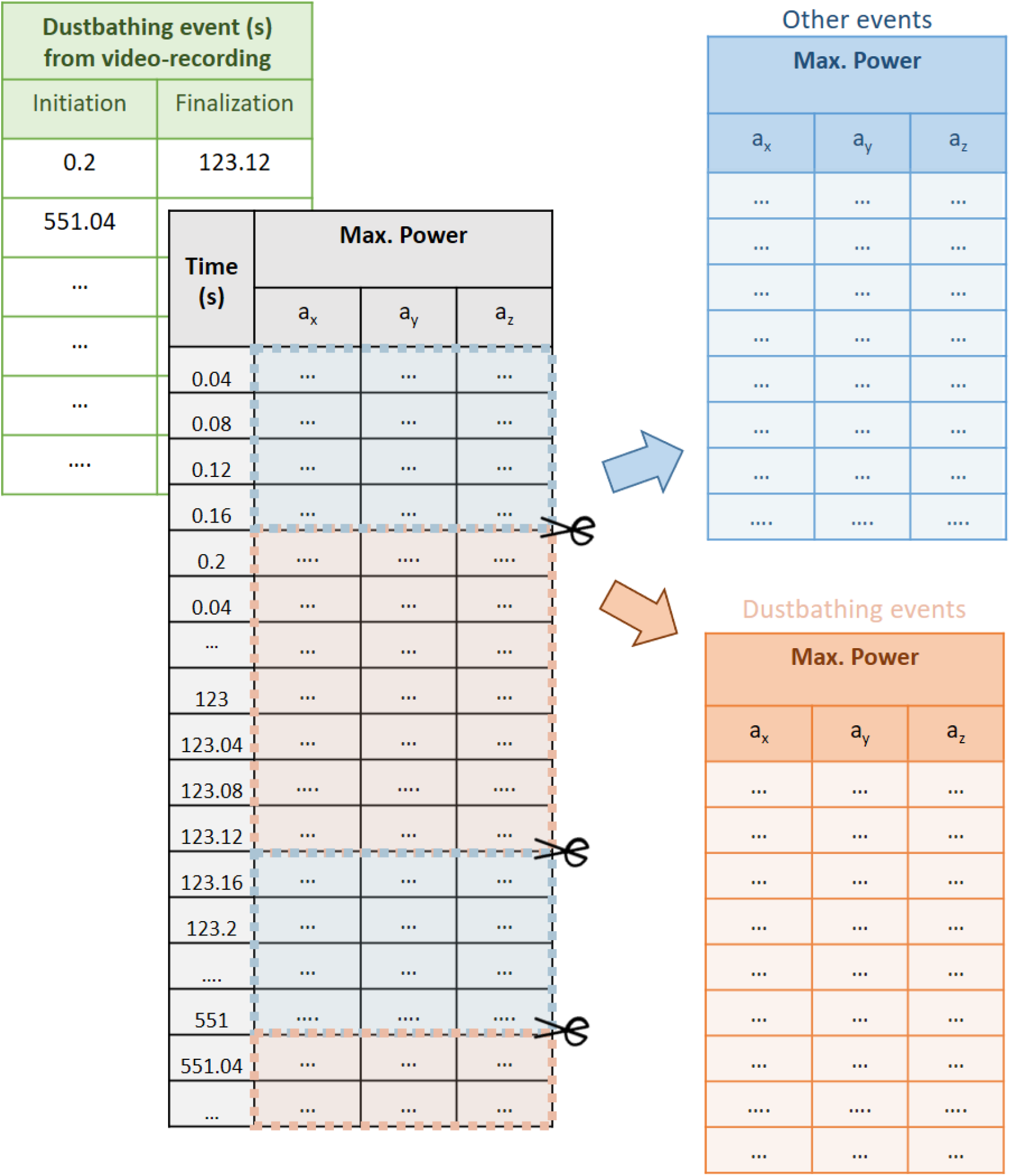
Schematic representation of segmentation process. Database of initiation and finalization of observed dustbathing events from video recordings (green table) was used to segment the database of the maximum power (i.e. squared modulus of wavelet coefficients) estimated with wavelet analysis of the three acceleration vectors (a_x_, a_y_ and a_z_; gray table), to create the two tables observed on the right corresponding to all segments without and with dustbathing (light blue and orange tables, respectively).

**Figure A3.**
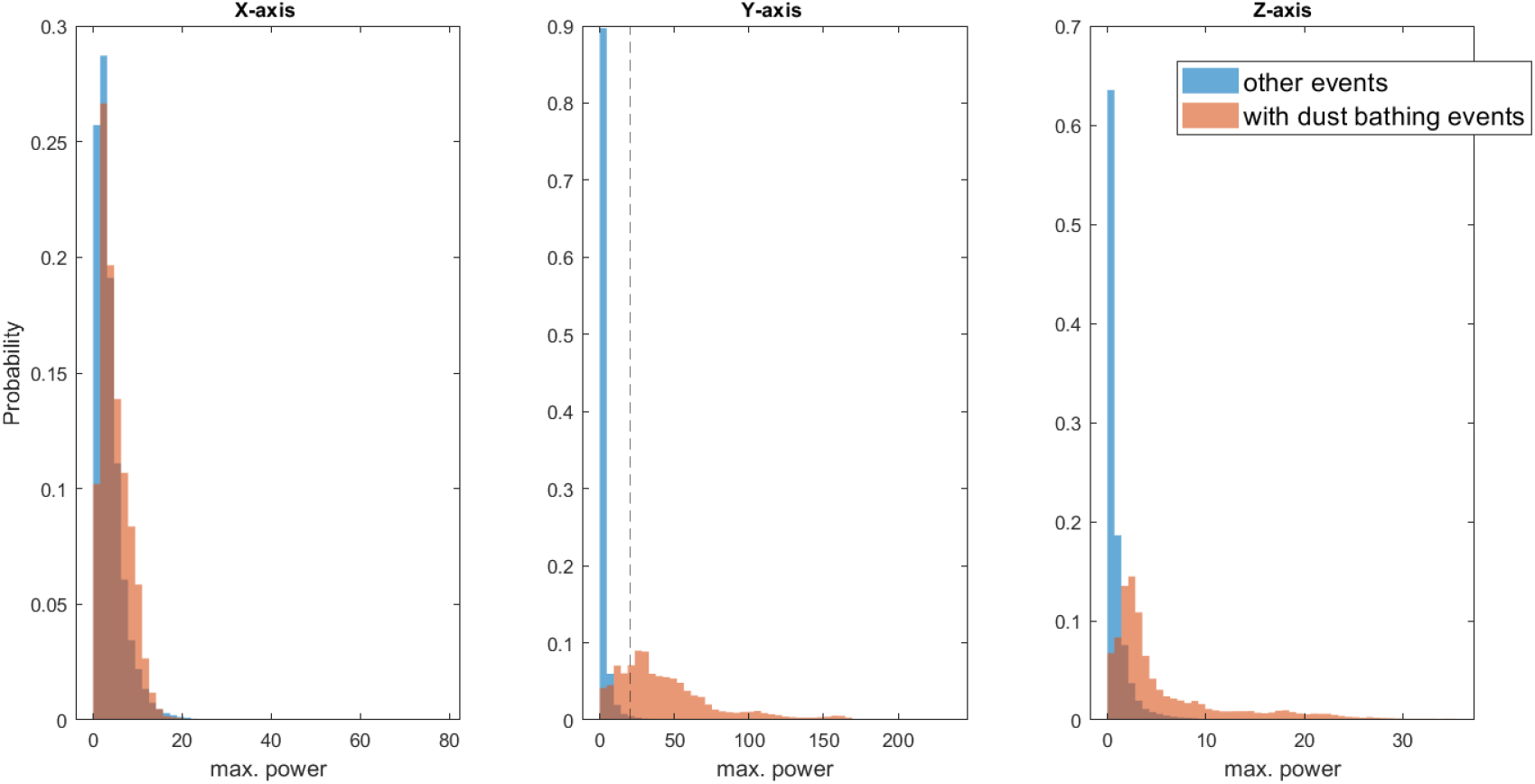
Probability distributions of the maximum power (i.e. squared modulus of the wavelet coefficients) estimated from wavelet analysis of the x-, y- and z-axis acceleration vectors from segments corresponding to periods without or with dustbathing as assessed from video-recordings. Dotted line marks the established threshold.

**Figure A4.**
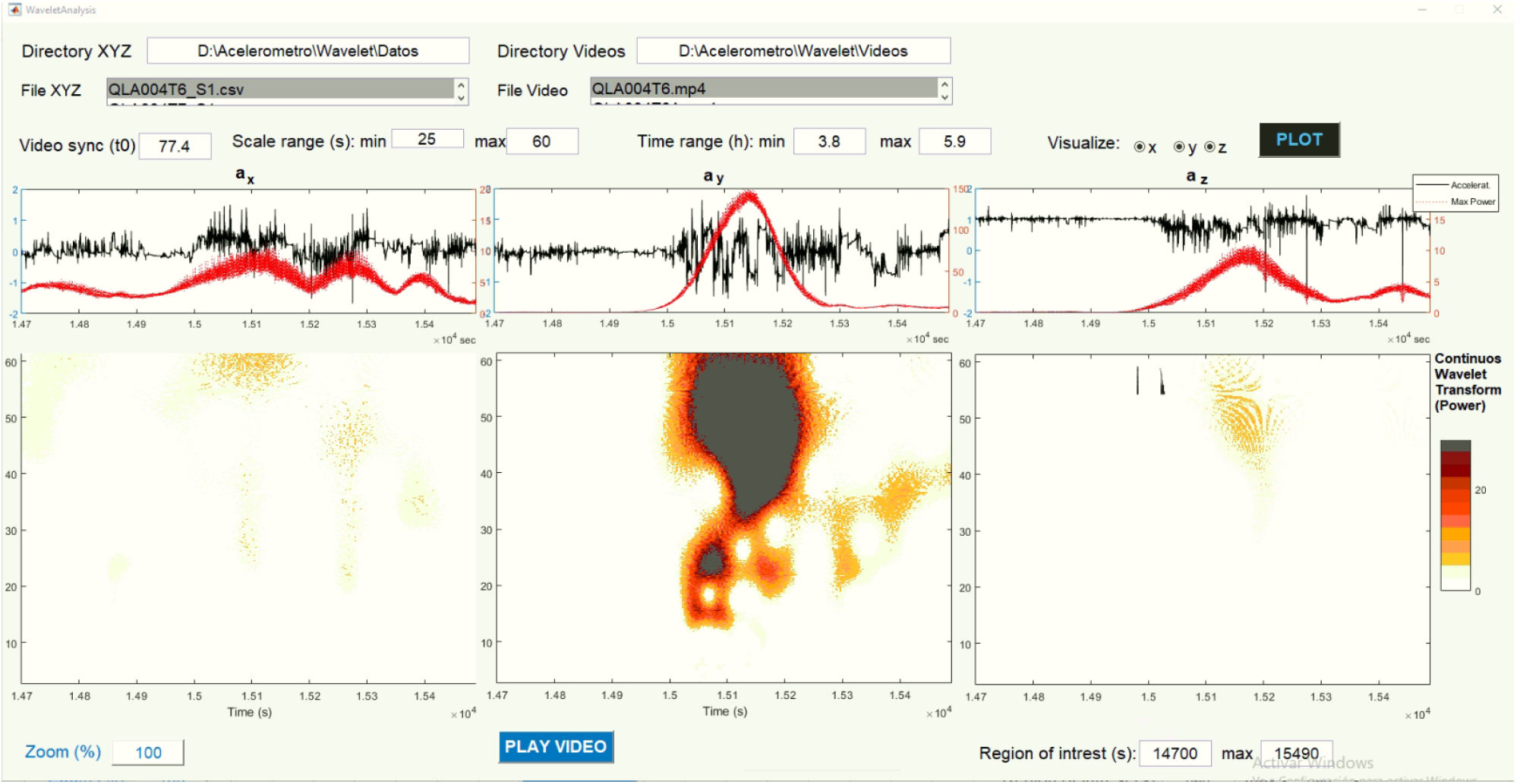
Zoom in on a temporal window with a dustbathing event in the costumed made MATLAB app. This is the same time series as shown in Figure 1C. Note the dark red area in the scpectrogram from the y-axis acceleration vector (middle panel) indicating large positive power values.

**Figure A5:**
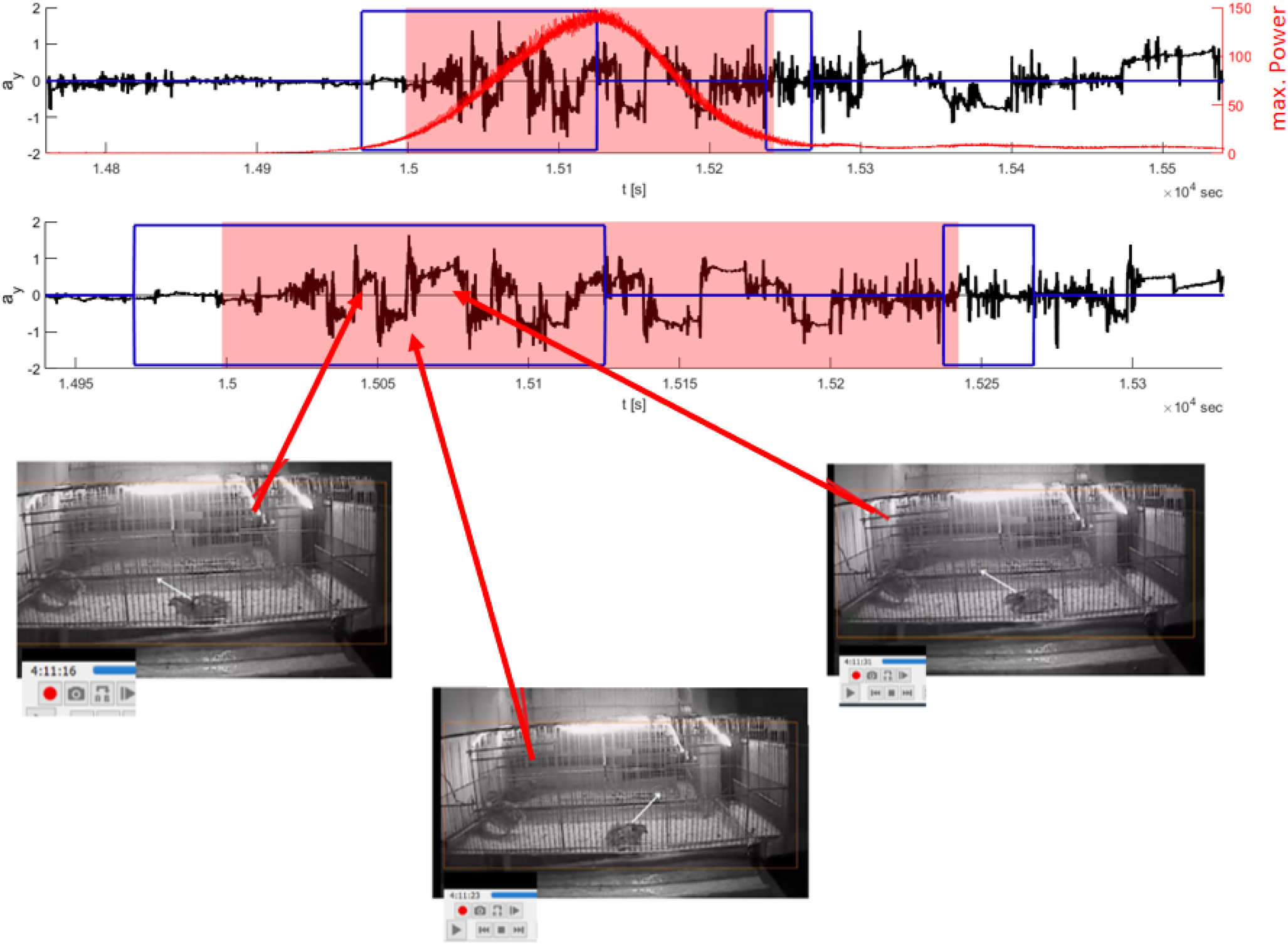
Example of dust bathing detection with wavelets. A) Comparison between y-axis acceleration vector (a_y_, black line) the estimated maximum power of the rhythm (red line) from the same example shown in Figure 1C, dust bathing event observed from video recordings (blue boxes), and automatic detection of dust baths by algorithm (red highlighted areas). B) Zoom in on the dust bathing event. Photographs show the association between the position of the bird’s body while lying down with peaks and valleys in the a_y_-rhythm. White arrows in photographs highlight the inclination of the bird’s body detected in the a_y_ -time series.

**Figure A6.**
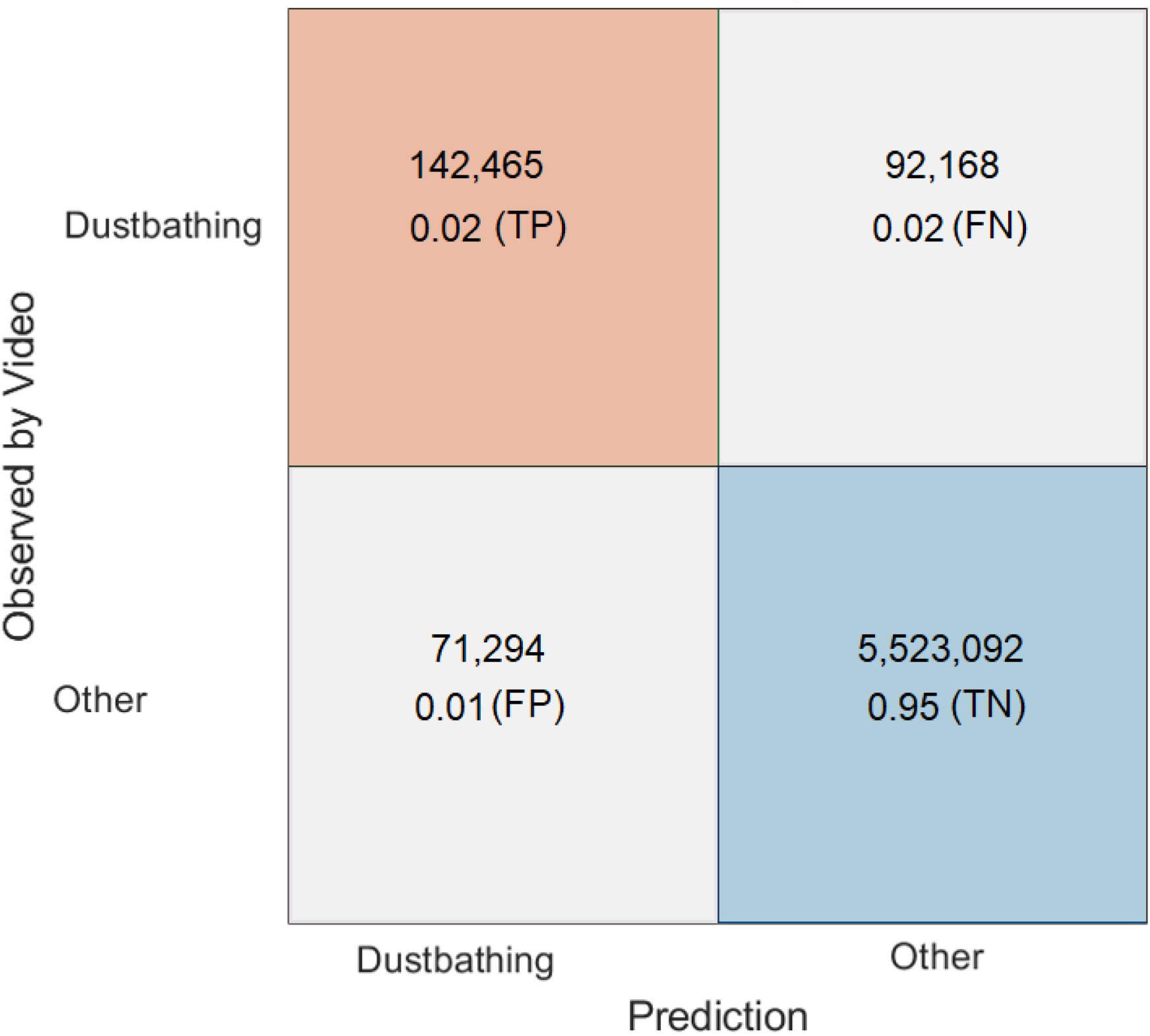
Overall confusion matrix detection of dust bathing focusing on data points. The number of data points from each class and the proportion of true-positives (TP), false-positive (FP), true-negative (TN) and false-negative (FN) rates are shown in squares. Please, compare to Figure 2C where the confusion matrix was performed for events.

**Video A1. Usage of the customized MATLAB application for visualizing acceleration vectors, wavelet scalograms and corresponding video-recording**.

**Video 2A. Visual comparison between acceleration vectors and video-recorded male behavior**. Note that two birds are present; only the male with the backpack was analyzed.

**Table A1.**
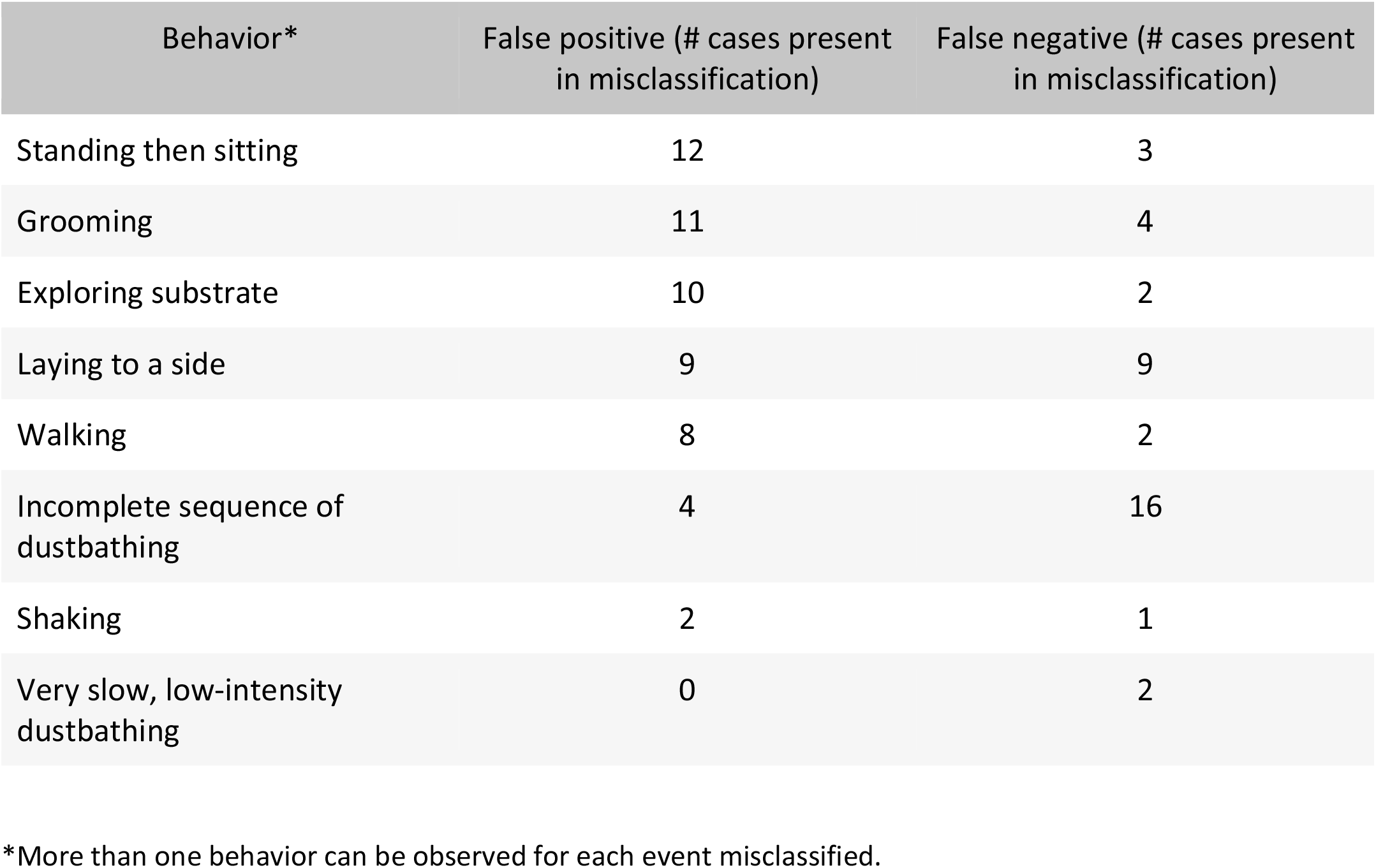
Behaviors observed in classification errors from video-recordings

| Behavior* | False positive (# cases present in misclassification) | False negative (# cases present in misclassification) |
| --- | --- | --- |
| Standing then sitting | 12 | 3 |
| Grooming | 11 | 4 |
| Exploring substrate | 10 | 2 |
| Laying to a side | 9 | 9 |
| Walking | 8 | 2 |
| Incomplete sequence of dustbathing | 4 | 16 |
| Shaking | 2 | 1 |
| Very slow, low-intensity dustbathing | 0 | 2 |

\*More than one behavior can be observed for each event misclassified.

